# Conformational switch in the alpha-synuclein C-terminus domain directs its fibril polymorphs

**DOI:** 10.1101/2022.02.10.479831

**Authors:** Cesar Aguirre, Yohei Miyanoiri, Masatomo So, Hajime Tamaki, Takahiro Maruno, Junko Doi, Nan Wang, Keiichi Yamaguchi, Kichitaro Nakajima, Yu Yamamori, Hiroko Inoura, Chi-Jing Choong, Keita Kakuda, Takahiro Ajiki, Yasuyoshi Kimura, Tatsuhiko Ozono, Kousuke Baba, Seiichi Nagano, Yoshitaka Nagai, Hirotsugu Ogi, Susumu Uchiyama, Yoh Matsuki, Kentaro Tomii, Yuji Goto, Kensuke Ikenaka, Hideki Mochizuki

**Affiliations:** Department of Neurology, Osaka University Graduate School of Medicine, 2-2 Yamadaoka, Suita, Osaka 565-0871, Japan; Laboratory for Ultra-High magnetic field NMR spectroscopy, Research Center for Next-Generation Protein Sciences, Institute for Protein Research, Osaka University, Japan, 3-2 Yamadaoka, Suita, Osaka 565-0871, Japan; Graduate School of Agriculture, Kyoto University, Kitashirakawa Oiwake-cho, Sakyo-ku, Kyoto 606-8502, Japan; Laboratory of Molecular Biophysics, Institute for Protein Research, Osaka University, Japan, 3-2 Yamadaoka, Suita, Osaka 565-0871, Japan; Department of Biotechnology, Graduate School of Engineering, Osaka University, Japan, 2-1 Yamadaoka, Suita, Osaka 565-0871, Japan; Graduate School of Engineering, Osaka University, Japan, 2-1 Yamadaoka, Suita, Osaka 565-0871, Japan; Artificial Intelligence Research Center (AIRC), National Institute of Advanced Industrial Science and Technology (AIST), 2-4-7 Aomi, Koto-ku, Tokyo, 135-0064, Japan; Department of Neurology, Faculty of Medicine, Kindai University, Osaka-Sayama, Osaka, Japan; Center for Quantum Information and Quantum Biology, Osaka University, Japan, 1-2 Machikane-yama, Toyonaka, Osaka 560-0043, Japan

**Keywords:** alpha synuclein, amyloid fibrils, fibril polymorphism, NMR

## Abstract

α-Synuclein (αSyn) inclusions are a pathological hallmark of several neurodegenerative disorders. While cryo-electron microscopy studies have revealed distinct fibril polymorphs across different synucleinopathies, the molecular switches controlling polymorphism remained unveiled. In this study, we found that fibril morphology is associated with the conformational state of monomeric αSyn. Through systematic manipulation of the ionic strength and temperature, we pinpoint two distinct polymorphs: a twisted morphology at low ionic strength and temperature, and a rod-like morphology at higher ionic strength and temperature. Most strikingly, we found that a specific conformational change in the C-terminal domain of the monomeric αSyn serves as the master switch for the formation of polymorphs. Interestingly, this conformational change can be triggered by calcium binding to the C-terminus, connecting environmental factors to specific fibril architectures. Our results unmask the C-terminal domain as a key player for orchestrating αSyn fibril morphology, providing significant insights into the fibrogenesis of αSyn.

**Significance Statement:** The αSyn C-terminus domain acts as the master switch programming its fibril polymorphism.

## Introduction

The accumulation of alpha-synuclein (αSyn) is a pathological hallmark of Parkinson’s disease (PD) and multiple system atrophy (MSA) (*1–3*). Although triggered by the same protein, these synucleinopathies exhibit distinctive clinical features. Previously, we have developed experimental strategies to study directly the structural characteristics of inclusions in patient’s brain employing Fourier-Transform Infrared microscopy (FTIR) and Small-Angle X-Ray Scattering (SAXS), proving that Lewy Bodies (LBs) contain cross β-sheet αSyn fibrils, while the amount of β-sheet structure in LBs and glial cell inclusions in MSA patients is different (*4–6*). More recently, direct observation by cryo-electron microscopy (Cryo-EM) of aggregates extracted from the brains of PD and MSA patients has revealed that these distinct clinical features map directly to specific fibril morphologies (*7*, *8*). While various strains of αSyn fibrils can be generated in vitro by manipulating the conditions of fibril formation (*9–12*), and some of them have been resolved by Cryo-EM (*13–16*), the fundamental question has persisted.

Despite these high-resolution structural snapshots of both in vitro and brain-extracted fibrils, the molecular mechanisms underlying the formation of polymorphs from a single protein remained one of the field’s most pressing mysteries. To address this central question, we asked: how do environmental parameters such as ionic strength and temperature reprogram the intrinsic structural properties of monomeric αSyn to generate distinct fibril architectures? While αSyn is canonically viewed as an intrinsically disordered protein (*17–19*), it can adopt partially folded conformations (*20–23*). Previous studies have suggested that these conformations could represent fibrillation intermediates (*22*, *24*, *25*), but their role in orchestrating the fibril polymorphism has remained elusive. Previous studies also established that ionic strength can modulate fibril polymorphs (*9–11*), and temperature influences the conformations of αSyn (*23*).

In this study, we describe a paradigm-shifting discovery: the C-terminal domain of monomeric αSyn functions as a molecular switch controlling fibril polymorphism. Through systematic manipulation of ionic strength and temperature, we generated and characterized two distinct fibril polymorphs: twisted and rod-like fibrils. Our results demonstrate that specific conformational changes in the monomeric αSyn C-terminus domain dictate the fibril architecture, while this particular domain remains outside the fibril core in the mature fibril (*13*, *14*). This breakthrough establishes also a link between early conformational events and the final fibril polymorphism. Although the fibril diversity studied in vitro is not directly indicative of pathological changes in humans, it points to the unexplored opportunities targeting the C-terminus domain to control polymorphism in the cellular environment.

## Results

### *α*Syn fibril morphologies can be controlled by fibrillation conditions

Previous studies hinted at ionic strength’s influence on fibril morphologies (*9–11*), but its molecular basis remained unclear. Thus, we launched a systematic investigation employing our HANABI instrument (*26–28*) to monitor fibril formation in real-time through thioflavin T (ThT) fluorescence. By precisely controlling conditions (50 mM TRIS pH 7.4, 0-500 mM NaCl and 37 °C), our result unveiled intriguing fibril growth patterns (**Figure 1.A**). Our analysis encompassing the full range of NaCl concentration revealed three distinct regimes of ThT intensity profiles: (i) consistently low maximal ThT intensities below 300 mM NaCl, (ii) a critical transition zone at 300 mM NaCl showing curves with both low and high maximal ThT, the inflection point, and (iii) exclusively high ThT intensities above 400 mM NaCl. This clear bifurcation became strikingly apparent when plotting the maximal ThT values as a function of NaCl concentration (**Figure 1.B**). Transmission electron microscopy (TEM) visualized the structural aspect of this phenomenon (**Figure 1.B**, upper and lower-right): low ThT signatures correspond to twisted fibrils, while high ThT intensities mark rod-like morphologies. Crucially, SDS-PAGE analysis confirmed near-complete fibrillation for both polymorphs (**Figure S1.A and B**), ruling out the possibility that yield variation as the source of ThT intensity differences.

**Figure 1.**
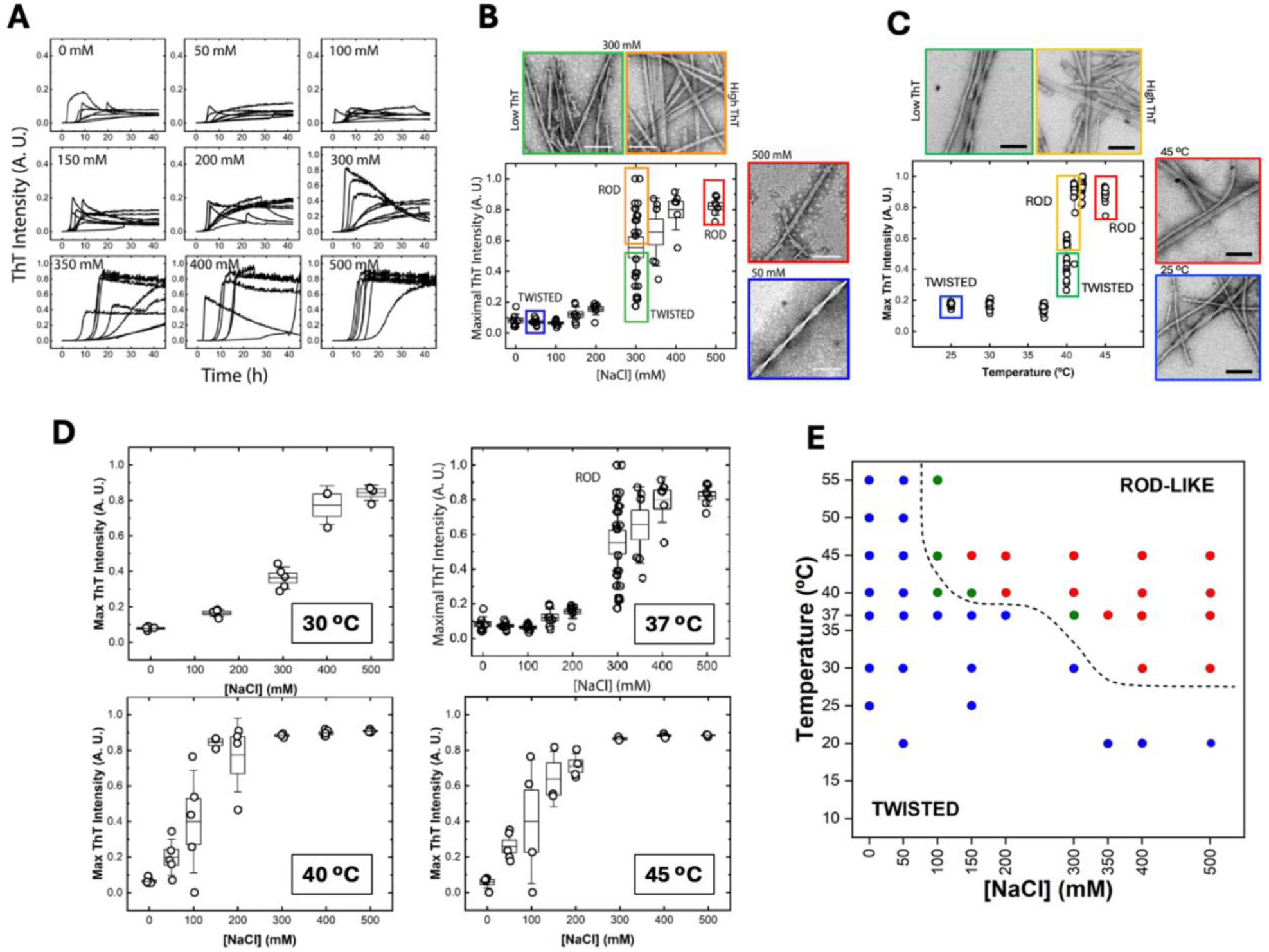
αSyn can generate rod-like and twisted polymorphs under different fibrillation reaction conditions. **A.** αSyn fibril formation kinetics monitored by ThT intensity at 37 °C, at the indicated concentrations of NaCl. **B.** Maximum ThT intensity values from the panel A plotted as a function of NaCl concentration. Boxes represent standard error of the mean; whiskers represent standard deviation. Also shown are the TEM images for selected fibrils, revealing: twisted fibrils at low maximal ThT intensity (blue and green squares), and rod-like fibrils at high maximal ThT intensity (yellow and red squares). White scale bar: 100 nm. **C.** Maximal ThT intensity values as a function of temperature an ionic strength of 0.15. Boxes represent standard error of the mean values; whiskers represent standard deviation. TEM imaging confirms formation of: twisted fibrils at low maximal ThT intensity (blue and green squares), and rod-like fibrils at high maximal ThT intensity (yellow and red squares). Black scale bar: 100 nm. **D.** Maximal ThT intensity values as a function of the NaCl concentration at different temperatures. Boxes represent standard error of the mean values; whiskers represent standard deviation. **E.** Phase diagram mapping fibril morphology as a function of NaCl concentration and temperature. Blue region indicates predominant twisted fibrils; green region corresponds to the inflection point where both rod and twisted fibrils coexist; red region shows predominant rod-like fibrils.

The role of temperature emerged as equally decisive. At the physiological ionic strength (0.15, fixed with NaCl), αSyn underwent a sharp conformational transition as temperature increased from 25 to 45 °C (**Figure 1.C**). Below 40 °C, fibrils exclusively exhibited low maximal ThT. At 40 °C we identified an inflection point, beyond which only the high ThT fibrils were observed. TEM analysis corroborated this dichotomy: the low ThT intensity fibrils invariably exhibited twisted morphology, while high ThT fibrils showed rod-like architectures, regardless of whether ionic strength or temperature drove their formation (**Figure 1.C**, upper and lower-right). Using SDS-PAGE, we also corroborated that, for both twisted and rod-like fibrils formed at different temperatures (**Figure S1.A**), the extent of the fibrillation reaction was nearly 100 % (**Figure S1.B**).

To put this finding into perspective, we determined a comprehensive polymorphism phase diagram mapping the interplay between the NaCl concentration and temperature, based on the maximal ThT values diagrams (**Figure 1.D)**. Remarkably, the inflection point for morphological transition shifts systematically towards lower NaCl concentrations as temperature rises, revealing a precisely controlled landscape of fibril polymorphism (**Figure 1.E)**. The map exposes a clear domain: twisted fibrils dominate at low temperature and low NaCl concentration (**Figure 1.E**, blue circles), while rod-like fibrils emerge under elevated conditions (**Figure 1.E**, red circles). These domains are separated by a sharp boundary, where both morphologies coexist (**Figure 1.E**, green circles).

These precise mapping reveals unprecedented molecular threshold governing fibril architecture. The existence of such clear phase boundaries suggests a specific molecular event, rather than random factors, that determines the formation of rod-like v.s. twisted fibrils. This discovery points to a fundamental mechanism controlling αSyn fibril polymorphism, demanding deeper investigation into the molecular basis of this intriguing phenomenon.

### Twisted and rod-like fibrils share a conserved core structure at the atomic level despite distinct overall architectures

To explore beyond morphological differences visible by TEM, we first performed proteinase K resistance (PKR) analysis as a molecular probe of fibril architecture (**Figure S1.C-D**). Over 60-minutes of proteolysis time course, we uncovered distinctive digestion profiles in the five highest molecular weight bands at the 30-min mark. Specifically, twisted fibrils, regardless of their formation at low NaCl or low temperature, exhibited characteristically weaker intensities in the second (B2) and fourth (B4) bands, together with a distinctly lower B4/B5 ratio compared to their rod-like counterparts. These result reveals fundamental differences in solvent-exposed regions between the two polymorphs.

For a deeper look at the molecular architecture of these fibrils, we turned to solid-state NMR spectroscopy. First, we generated uniformly ^13^C,^15^N-isotopically labeled αSyn fibrils under four critical conditions: i) twisted fibril at low NaCl concentration, ii) rod-like fibril at high NaCl concentration iii) twisted fibril at low temperature, and iv) rod-like fibril at high temperature. Each polymorph’s identity was rigorously confirmed through both TEM visualization and PKR analysis (**Figure 2.A**). Most remarkably, our comprehensive structural analysis reveals an unexpected puzzle: while the fibril surfaces display distinct proteinase K accessibility patterns unique to each polymorph, their core structures remain strikingly similar across all conditions, as evidenced by their ssNMR signatures (**Figure 2.B**). This finding aligns with, yet substantially extends, previous cryo-EM studies (*13*, *15*) showing that both twisted and rod-like polymorphs comprise two protofilaments sharing a highly conserved core. The key distinction between polymorphs appears to arise not from fundamental differences in their core structure, but rather from specific variations in protofilament interface interactions. This is together suggesting that while the basic building block of αSyn fibrils remain constant, the final architecture is given by precise molecular interactions that guide protofilament assembly. This insight raises a crucial question of what molecular mechanisms control these assembly patterns at the earliest stages of fibril formation.

**Figure 2.**
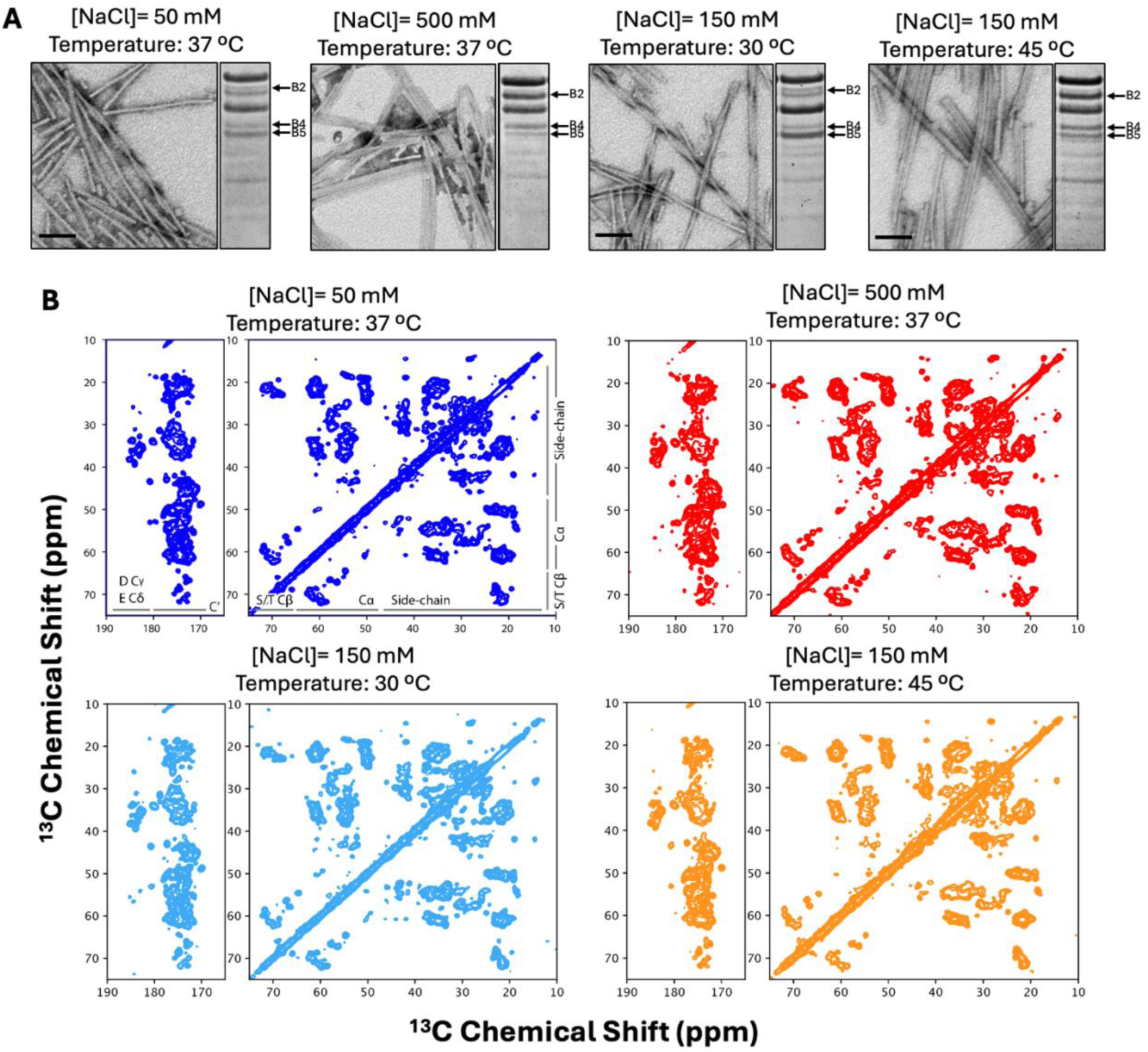
Solid-state NMR analysis of twisted and rod-like fibrils. **A.** Twisted and rod-like fibrils were generated at the NaCl concentration and temperature conditions indicated in the upper part of the panels. The morphological properties of the fibrils were confirmed by TEM visualization (scale bar: 100 nm) and PKR after 30 min of digestion (arrows: distinctive bands of each polymorph). **B.** 2D ^13^C-^13^C DARR spectra of twisted and rod-like fibrils produced under the specified conditions. Aliphatic-carbonyl and aliphatic-aliphatic ^13^C correlations are shown in the left and right panels, respectively, for each sample. Intra-residue correlations spanning 1-3 bond were observed with a short ^13^C-^13^C DARR mixing time of 20 ms. No apparent differences among all spectra.

### The C-terminus domain master switch correlates with fibril polymorphism

To answer this question, we turned to a radical hypothesis: could fibril polymorphism be predetermined at the monomer state? Our first clue came from size exclusion chromatography (SEC) analysis of monomeric αSyn (**Figure S2.A,** upper). The full-length αSyn monomer (14.6 kDa) was observed not as a sharp peak, but as an asymmetric distribution spanning apparent molecular weights of 15-60 kDa (for hypothetical spherical-shaped globular proteins), with its center around 50 kDa. Crucially, Western Blot analysis confirmed that every fraction contained exclusively full-length monomers (**Figure S2.A**, lower), revealing this broad distribution as a signature of distinct conformation states rather than oligomerization.

This conformational landscape proved exquisitely sensitive to ionic strength. As NaCl concentrations increased from 0 to 500 mM, we observed a continuous increase in the retention volume, reaching a plateau above 300 mM NaCl (**Figures S2.B**). Analytical ultracentrifugation (AUC) reinforced this finding, corroborating the monomeric state of αSyn in solution and revealing a decrease in the sedimentation coefficient (*s*_w_) with increasing NaCl concentration (**Figure S2.C-D and Table S1**). Together, these results indicate that conformations yielding low retention volumes and high *s*_w_ values, characteristic of low ionic strength, generate twisted fibrils, whereas those at the retention volume plateau region with low *s*_w_ values invariably produce rod-like fibrils.

Temperature’s influence proved equally dramatic, though technically more challenging to map. While SEC analysis revealed a linear correlation between retention volume and temperature (**Error! Reference source not found.**), the convolution of the effect of temperature on both αSyn conformations and the SEC column properties precluded detailed analysis on temperature-dependent conformational effects through this technique alone.

To unmask the precise nature of these polymorphism-determining conformations, we deployed ^1^H-^15^N HSQC NMR spectroscopy across our phase diagram, following four strategic routes indicated in **Figure 3.A**. Using the spectrum obtained at 0 mM NaCl and 10 °C as the reference point (**Figure 3.A,** marked with a cross), we revealed dramatic changes in the peak intensities across all routes (**Figure 3.B**), while not in the chemical shifts (**Error! Reference source not found..B**). Most strikingly, we identified a critical region spanning residues 110-130 in the C-terminal domain that displayed unique behavior. This region showed distinctive pattern in peak intensity changes across each route (highlighted in dark purple, red, dark yellow and blue for routes R1-R4, respectively). While the N-terminus domain and NAC region showed general intensity reductions with increasing temperature or NaCl, this C-terminal region exhibited route-specific responses that correlated with fibril morphology: in route R1, a slow intensity decrease with temperature increase; in route R2, a more rapid decrease, resulting in very small intensities (< 10% of the reference) at above 40 °C; in route R3, a more gradual reduction that leveled off at around 300 mM NaCl; finally, in route R4, rapid decrease resulting in <10 % intensity above 300 mM NaCl.

**Figure 3.**
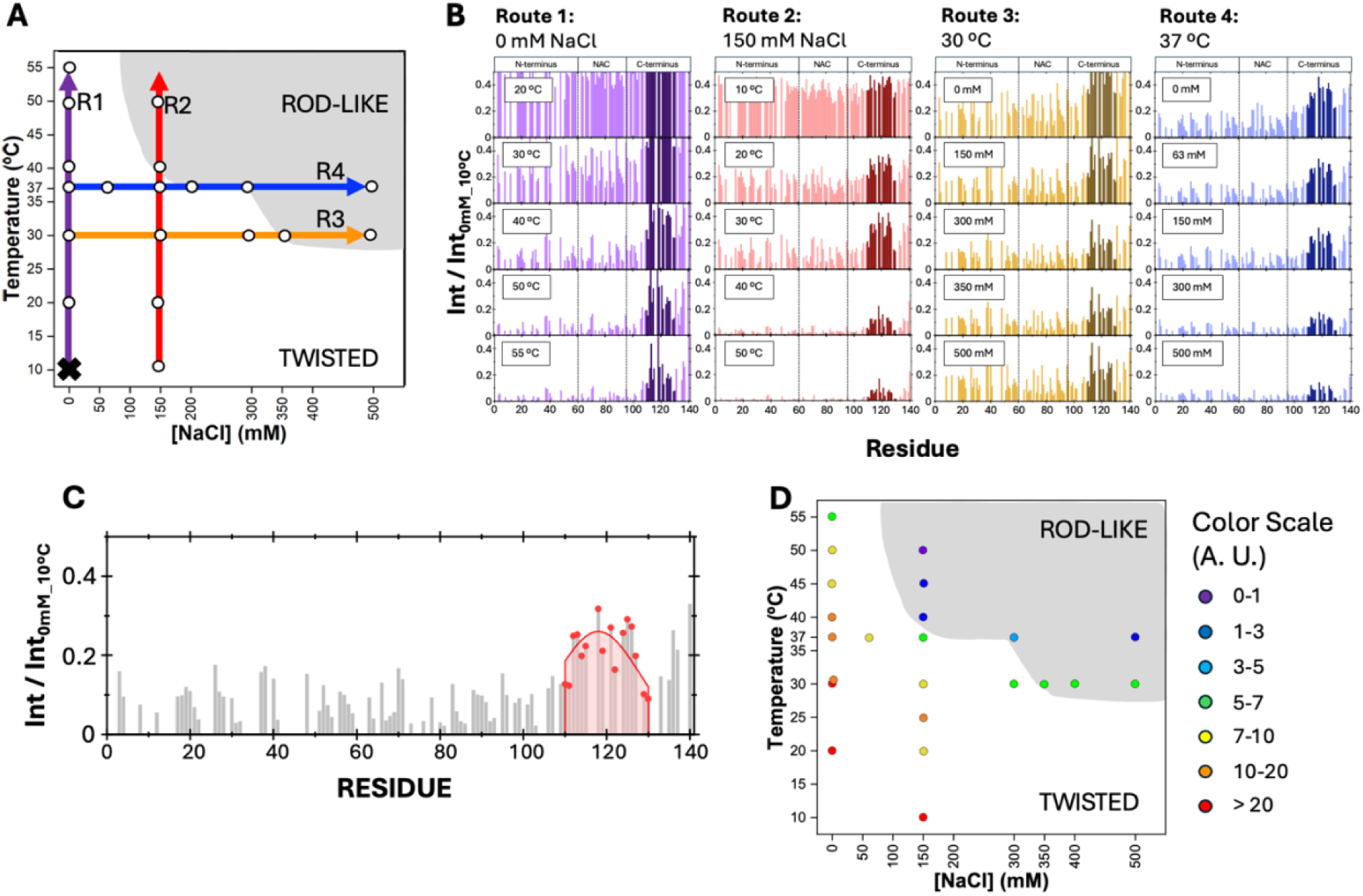
Solution NMR analysis of αSyn monomer conformational states. **A.** Schematic representation of four experimental routes examining the monomeric αSyn conformational changes, using ^1^H-^15^N HSQC measurements under polymorph-inducing conditions. The reference spectrum was taken at 10 °C without NaCl (black cross). R1: Temperature variation (10 °C to 55 °C) at 0 mM NaCl; R2: Temperature variation (10 °C to 55 °C) at 150 mM NaCl; R3: NaCl concentration variation (0 to 500 mM) at 30 °C; and R4: NaCl variation (0 to 500 mM) at 37 °C. White circles represent the tested conditions. **B.** Peak intensity ratio observed in the ^1^H-^15^N HSQC spectra at selected points along routes R1 to R4, normalized to the reference spectrum (no NaCl at 10 °C). Peaks corresponding to the C-terminus residues (residues 110-130) are highlighted across all routes. **C.** Quantification of the overall peak intensity for residues 110-130 through Gaussian curve fitting and area integration. **D.** Integrated peak intensities (colored circles) from panel C mapped onto the fibril morphology phase diagram.

Integration of the peak intensity ratio (**Figure 3.C)** allowed us to map these conformational changes directly onto our morphology phase diagram (**Figure 3.D**). The results revealed an unprecedented molecular switch: when the peak intensity ratio for residues 110-130 fall below 3 arbitrary units, a rod-like polymorph emerges; above 7 units, twisted fibrils dominate. This sharp threshold represents a specific conformational transition in the C-terminus domain that determines fibrils morphology. Crucially, we ruled out oligomerization effects through immediate post-NMR SEC analysis (**Figure S5.A**) and confirmed the persistence of disordered structure through far-UV CD spectroscopy (**Figure S5.B**). These results indicate that the conformational switch occurs within a still-disordered ensemble, consistent with previously reported partially folded conformations (*23*), but now related to polymorphism control.

### Ca^2+^ binding to the C-terminal domain induces rod-like fibrils formation

We also discovered calcium binding to the C-terminal domain specifically drives the formation of the rod-like fibril. While previous studies showed that calcium binds αSyn and accelerates fibrillation (*29*, *30*), our findings reveal unprecedented role of calcium in determining fibril morphology under physiological conditions. ^1^H-^15^N HSQC spectra recorded at physiological ionic strength (0.15, 37 °C, with and without Ca^2+^, **Error! Reference source not found..A**) (see Materials and methods) unveiled a clear calcium-induced conformational change. Most notably, residues 110-130 in the C-terminal domain showed a dramatic decrease in peak intensities upon calcium binding (**Figure 4.A**), while chemical shift patterns remained largely unchanged (**Error! Reference source not found..B**).

**Figure 4.**
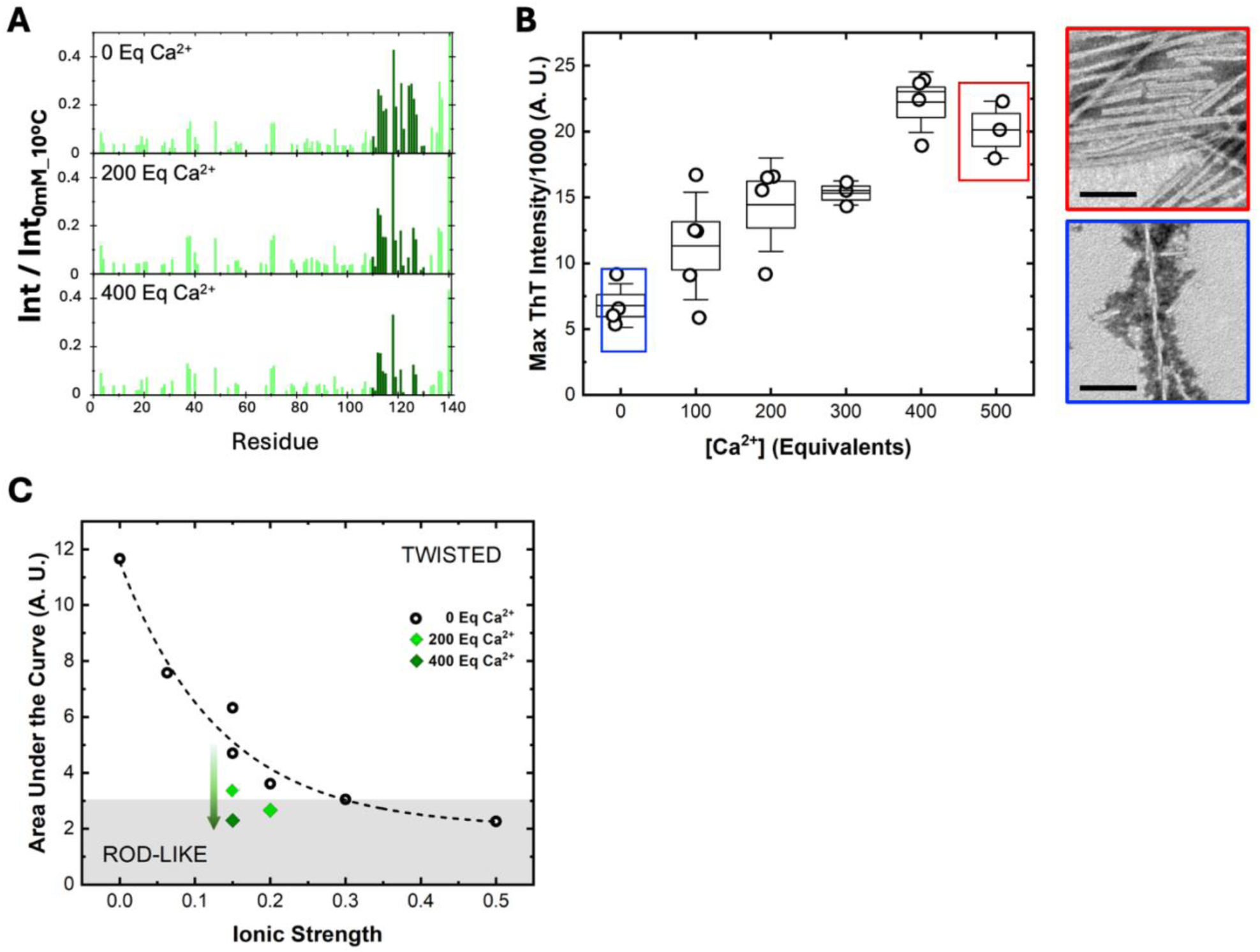
Calcium-induced formation of rod-like fibrils under modest ionic strength conditions. **A.** Peak intensity ratio from the ^1^H-^15^N HSQC spectra at varying Ca^2+^ concentrations but constant physiological ionic strength (0.15), normalized to the reference spectrum taken without no NaCl nor Ca^2+^ at 10 °C. C-terminus residues (110–130) showed a characteristic local peak pattern seen in Figure 3.B. **B.** Box-and-whisker plot showing maximal ThT intensities from fibrillation reactions, at constant ionic strength of 0.15 across different Ca^2+^ concentrations. Boxes indicate standard error of the mean; whiskers represent standard deviation. TEM images demonstrate the morphology difference: low maximal ThT intensity condition produced twisted fibrils (blue square), while high maximal ThT intensity conditions yielded rod-like fibrils (red square). Black scale bar: 100 nm. **C.** Integrated peak intensities for residues 110-130 mapped onto the morphology phase diagram for calcium-containing samples (green diamonds) at ionic strength of 0.15 or 0.20, and calcium-free samples at different ionic strength (black open circles, same values as those shown in Figure 3**.D**), at 37 °C. Calcium addition induces a conformational shift favoring rod-like over twisted fibril formation.

This conformational change translated into a profound effect on fibril morphology. By systematically varying Ca^2+^ concentration at physiological ionic strength (0.15), we observed a clear transformation in fibril architecture. TEM analysis revealed an exclusive population of twisted fibrils in calcium free condition, while addition of calcium (500 Eq.) triggered formation of rod-like fibrils (**Figure 4.B**, right). This morphological switch was accompanied by a systematic increase in ThT fluorescence, reaching saturation above 400 Eq. Ca^2+^ (**Figure 4.B**, left). SDS-PAGE analysis confirmed near-complete fibrillation under both conditions (Error! Reference source not found.**.B**).

The Ca^2+^-induced rod-like fibrils exhibited a unique structural signature in PKR digestion assays (Error! Reference source not found.**.C**). While Ca^2+^-free conditions generated the characteristic twisted fibrils digestion pattern, calcium-containing samples showed a unique profile distinct with persistently faint B2 and B4 bands. These results indicate that calcium binding induces rod-like fibrils with a novel arrangement of solvent-exposed regions, distinct from those formed under high NaCl or high temperature conditions.

Quantitative analysis of the residues 110-130, in terms of the area under the curve for the peak intensities, revealed that calcium binding reduces the peak intensity ratios to levels matching those observed under high ionic strength conditions generating rod-like fibrils (**Figure 4.C,** comparing green diamonds with black open circles). This comparison demonstrates that, at physiological ionic strength, where twisted fibril is the predominant polymorph, calcium binding reconfigures the C-terminal domain to mirror conformational states achieved by high NaCl conditions, thereby directing the formation of rod-like fibrils.

## Discussion

Our systematic investigation of αSyn fibril polymorphism in vitro has uncovered a fundamental principle that the conformational state of monomeric αSyn, specifically its C-terminal domain, switches the final fibril architecture. By precisely controlling NaCl concentration and temperature in fibrillation reactions, we constructed a comprehensive phase diagram that revealed two predominant fibril polymorphs: twisted fibrils that forms under low NaCl concentration / temperature conditions and rod-like fibrils emerging at higher values. This phase diagram proved instrumental in unveiling the molecular mechanisms underlying polymorphism.

While twisted and rod-like fibrils exhibit distinct morphological features and unique proteinase K digestion patterns, our ssNMR experiments revealed a surprising conservation of their core kernel structure. This finding aligns with previous cryo-EM studies (*13*, *15*), showing that both polymorphs share similar protofilaments cores, with morphological differences arising from distinct protofilament interfaces, affected probably by the C-terminus conformational state. However, our study goes beyond morphological characterization to reveal the molecular origins of these differences.

A key breakthrough emerged from our analysis of monomeric αSyn conformations in solution. Although αSyn is traditionally viewed as an intrinsically disordered protein, our SEC experiments revealed that its conformational ensemble is not random but responds systematically to environmental conditions. The conformational transitions observed through changes in retention volume reached a critical plateau at 300 mM NaCl, precisely matching the inflection point in the maximal ThT intensity where fibril morphology switched from twisted to rod-like. This remarkable correlation provided the first evidence that specific monomeric conformations are related with the final fibril architecture.

Solution ^1^H-^15^N HSQC measurements provided insights about the molecular mechanism behind this phenomenon, revealing a previously unrecognized regulatory role of the C-terminus domain. We identified a critical region spanning residues 110-130 that acts as a molecular switch for fibril polymorphism. While both the N-terminus domain and the NAC region showed general responses to changing conditions, only the C-terminus exhibited behavior that precisely correlated with fibril morphology. This finding rules out N-terminal and NAC regions as determinants of polymorphism, focusing attention on the C-terminus as the master regulator of the fibril architecture.

The importance of this discovery became evident when we quantified the C-terminal conformational state using area-under-curve analysis of peak intensities. We identified a critical threshold value of ∼3 A. U. that marks the transition from twisted to rod-like fibril formation. This empirical value possibly represents a specific conformational state of the C-terminus domain that enables rod-like polymorph formation. Previous studies have noted that C-terminal modifications can affect fibrillation kinetics through various mechanisms, including calcium binding (*30*) and proline isomerization (*31*, *32*). Uversky *et al* (*23*) reported temperature-induced partial folding of αSyn, but the link between these conformational changes and fibril polymorphism remained undiscovered until now.

Our calcium-binding experiments provided compelling validation of this mechanism. Calcium ions induced identical changes in the C-terminal region as high ionic strength and temperature, driving rod-like fibril formation even under otherwise twisted-favoring conditions. This demonstrates that C-terminus conformation, rather than bulk solution properties, seems to control fibril architecture. Although we cannot exclude the possibility that other divalent ions that bind the C-terminus domain (*33*) have a similar effect than calcium, our findings establish calcium, whose dysregulation has been proven to be associated to the development of neurodegenerative diseases (*34*), as a master regulator of αSyn fibril polymorphism through its specific interaction with the C-terminal domain, providing a crucial link between cellular homeostasis and pathological αSyn aggregation patterns. Future studies should further characterize conformational states of the C-terminus domain in function of the environmental factors based on, for example, solvent protection properties measured by the H-D exchange NMR experiments.

Based on these findings, we propose a model for understanding αSyn aggregation as illustrated in **Figure 5**. The C-terminal domain serves as a conformational switch: when residues 110-130 adopt a partially folded state, whether induced by salt, temperature, or calcium, rod-like fibrils form. In contrast, when this region remains flexible, twisted fibrils emerge. While a specific mechanism of how the C-terminus conformation guides the fibrillation process to generate specific polymorphs is to be answered, we propose that, since the protofilament core structure remains conserved, as shown by our ssNMR data, the C-terminus conformation influences the crucial protofilament interfaces that determine final fibril architecture. Experiments that identify interaction sites between the C-terminal domain and the fibril core during the fibril formation using, for example, paramagnetic relaxation enhancement (PRE) ssNMR measurements could further clarify this mechanism.

**Figure 5.**
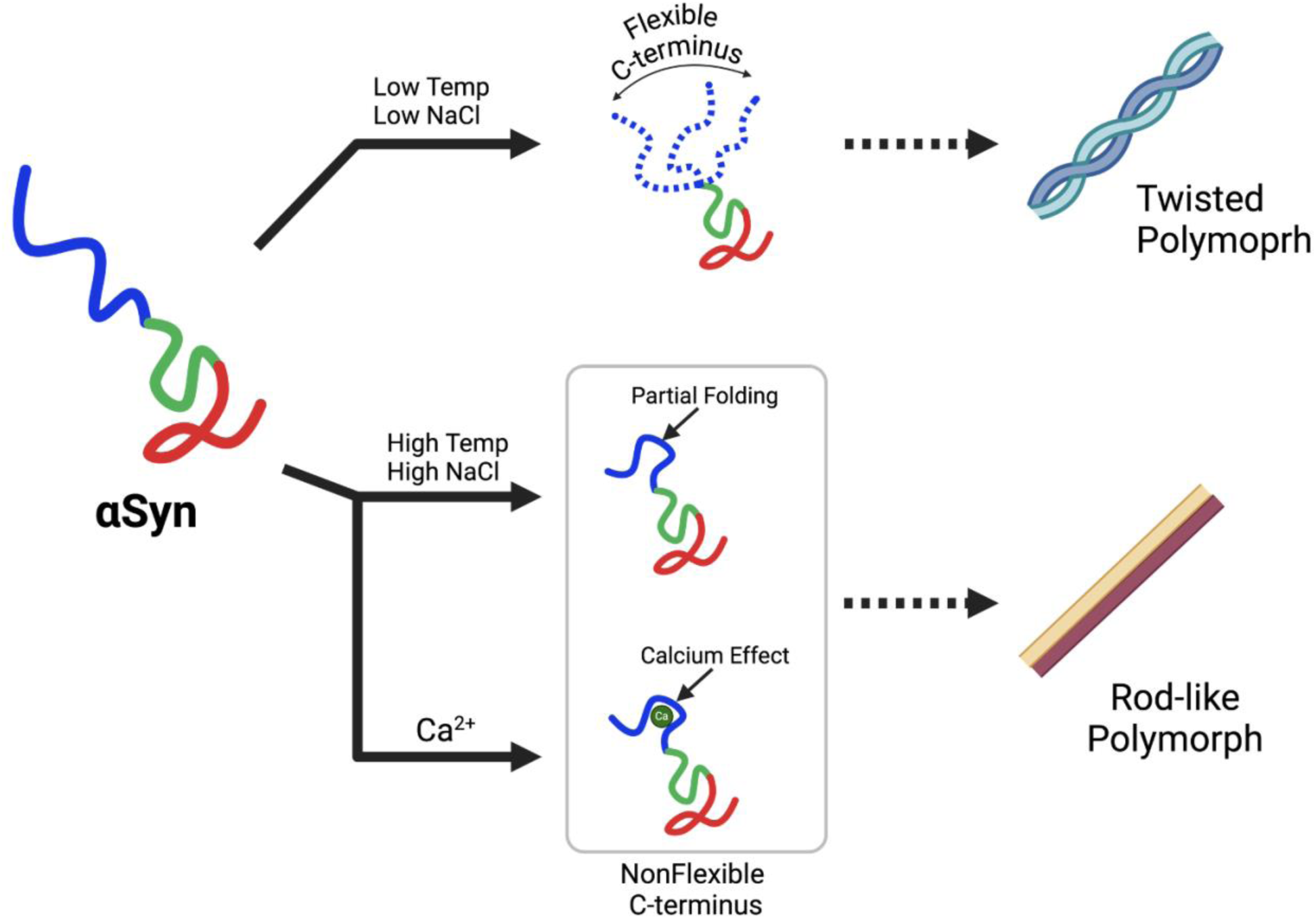
Relationship between αSyn C-terminus dynamics and fibril morphology. The conformational state of the C-terminal domain determines the fibril polymorphism: high C-terminus flexibility promotes twisted fibrils, while partial C-terminal folding drives the assembly of rod-like fibrils. Created with BioRender.com

This model has profound implications for understanding pathological αSyn aggregation. While we focused on ionic strength and temperature as experimental tools, our findings suggest that any cellular factor affecting C-terminal conformation, like binding partners, such as lipids (*12*, *35*, *36*), or local chemical environments, could influence fibril polymorphism to generate the polymorphs observed in fibrils extracted from the brain of patients (*7*, *8*). Indeed, this notion establishes a link between recently observed differential lysosomal damages and fibril morphologies, that results in distinct pathologies (*37*). Identifying the C-terminus domain as a key factor is particularly remarkable as the C-terminal domain undergoes in the complex environment of the brain, where it is target of numerous physiological modifications, such as phosphorylation, ubiquitination, truncation, metal coordination, etcetera (*38*), each potentially directing specific fibril morphologies through a mechanism similar to the model that we propose. In the future, this insight might enhance our understanding of the complex cellular events that trigger pathological aggregation and opens new possibilities: modulating C-terminal conformation could prevent formation of pathological polymorphs at their source.

Moreover, our discovery that domains outside the fibril core can regulate aggregation may represent a general principle in protein aggregation diseases. The C-terminus domain’s role in directing fibril architecture demonstrates how regions excluded from the final fibril structure can nevertheless determine its fate. This paradigm shift suggests new approaches to understanding and treating synucleinopathies by focusing on the conformational state of monomeric protein rather than mature fibrils.

## Materials and Methods

All chemicals and reactants were obtained from Sigma-Aldrich or Nacalai Tesque (Kyoto, Japan). All reagents were used without further purification. Water was purified to a resistivity of 18 MΩ cm^-1^ using a Millipore Gradient deionizing system.

### Protein purification

Human WT αSyn was purified from *Escherichia coli* as described previously (*39*). Briefly, a plasmid containing WT human αSyn was expressed in *E. coli* BL21 (DE3) cells (Novagen, Merck, San Diego, CA, USA). The cells were suspended in purification buffer (50 mM TRIS-HCl pH7.4, 1 mM EDTA, 0.1 mM DTT and 0.1 mM PMSF), disrupted by sonication, and centrifuged. Streptomycin sulfate (final 2.5% w/w) was added to the supernatant, and centrifugation was repeated. The supernatant was heated to 85 °C in a water bath and centrifuged. The supernatant was precipitated by the addition of solid ammonium sulfate to 70% saturation, centrifuged, dialyzed overnight, applied to a Resource-Q column (GE Healthcare, Little Chalfont, UK) coupled to an AKTA Explorer HPLC Instrument (Amersham Biosciences), and eluted with a linear gradient of 0–1 M NaCl. αSyn-enriched fractions (as determined by sodium dodecyl sulfate-polyacrylamide gel electrophoresis SDS-PAGE/Coomassie blue staining) were pooled and further purified by size exclusion chromatography (SEC) using a Superdex 200 26/600 PG column (GE Healthcare, Little Chalfont, UK) equilibrated with 50 mM TRIS-HCl pH 7.4 supplemented with 150 mM NaCl. The fractions containing αSyn (as determined by SDS-PAGE/Coomasie blue staining) were joint, dialyzed versus deionized water, acidified with 10 mM HCl and loaded onto a Reverse Phase Cosmosil Protein R x 250 mm Preparative Column (Nacalai-Tesque, Kyoto, Japan) coupled to a Gilson HPLC Unipoint System Instrument (Gilson, USA), and eluted with a linear gradient of 30 % - 90 % acetonitrile. The pure fractions were combined and flash-frozen in liquid nitrogen, lyophilized and stored at -80 °C until use. The protein purity was confirmed to be greater than 95% by SDS-PAGE and matrix-assisted laser desorption/ionization mass spectrometry. The protein concentration was determined by UV absorption at 280 nm using an extinction coefficient ε_1%_ of 3.54 (*40*).

### Kinetics of fibril formation followed by thioflavin T (ThT)

The formation of αSyn amyloid fibrils was studied by preparing solutions of αSyn (0.5 mg mL^-1^) in fibrillation buffer (50 mM TRIS-HCl pH 7.4), containing 10 μM ThT in a total volume of 200 µL, which were placed in each well of a 96-well microplate, and run under the conditions of ionic strength and temperature indicated in the text. The ThT fluorescence intensity was monitored using a HANdai Amyloid Burst Inducer (HANABI-2000) equipment, developed by our group in collaboration with CORONA ELECTRIC, Ibaraki, Japan, consisting of a transducer driving system, an acoustic-intensity measurement system, and a fluorescence measure system, in which a microplate reader was combined with a multichannel ultrasonication system that applies ultrasonic irradiation independently to each individual well of the microplate via a miniaturized ultrasonic resonator attached to each well. Ultrasonication was applied to accelerate amyloid formation at an optimized frequency of 30 kHz (*26–28*), in cycles of 300 ms of irradiation and 500 ms of quiescence. The kinetic parameters of fibril formation were determined by fitting the experimental data to empirical equations (*41*).

### Transmission electron microscopy (TEM)

TEM imaging was achieved on a Hitachi H-7650 transmission electron microscope (Hitachi, Tokyo) operated at 80 kV. Samples were diluted 1:1 with deionized water, and 10 µL of this solution was placed onto copper grids (400-mesh) covered with carbon-coated collodion film (Nisshin EM, Tokyo) and incubated for 1 min at room temperature. The samples were negatively stained with 10 µL of a 1% (w/v) solution of phosphotungstic acid (PTA), incubated for 1 min, and finally washed again with 10 µL of deionized water. The magnification working interval ranged from 5 000× to 20 000×.

### Proteinase K resistance (PKR) assay

αSyn fibrils (0.5 mg mL^-1^) in fibrillation buffer were digested using proteinase K (1 µg mL^-1^, Roche) at 37 °C and 400 rpm for different time intervals. To stop the reaction, the samples were incubated for 5 min at 95 °C, mixed with loading buffer (50 mM TRIS-HCl, pH 6.8, 4% SDS, 2% β-mercaptoethanol, 12% glycerol and 0.01% bromophenol blue) and incubated at 95 °C for additional 10 min. The digestion patterns were analyzed by SDS-PAGE followed by Coomassie Brilliant Blue staining. The first five products of digestion, namely B1 to B5, were employed for the analysis.

### Analytical size-exclusion chromatography (SEC)

Solutions of αSyn at a concentration of 1 mg mL^-1^ were filtered through a 0.22 µm size-pore membrane and loaded onto a 75 Increase 10/300 GL SEC column (GE Healthcare). The flow rate was set to 0.5 mL min^-1^. The protein absorption at 220, 250 and 280 nm was monitored.

### Analytical Ultracentrifugation (AUC)

Sedimentation velocity (SV) experiments were performed at 20 °C using an Optima AUC analytical ultracentrifuge (Beckman Coulter) equipped with an An-60Ti rotor using 12-mm double-sector aluminum centerpiece with sapphire windows. αSyn was dissolved in the buffer (50 mM TRIS-HCl pH 7.4; 50 mM NaCl, 50 mM TRIS-HCl pH 7.4, 150 mM NaCl; and 50 mM TRIS-HCl pH 7.4, 500 mM NaCl). The protein concentrations were adjusted to absorbance values of 0.8 at 230 nm in the working buffer at 1-cm path length. 390 µL of prepared sample solution and 400 µL of buffer were loaded to the appropriate sector of centerpiece. Sedimentation data were collected every 2 min at 60,000 rpm with a radial increment of 0.001 cm using absorbance optics. The detection wavelength was set at 230 nm.

The distribution of the sedimentation coefficient was determined using the continuous *c*(*s*) distribution model in the program SEDFIT (*42*). The range of the sedimentation coefficients for fitting was 0-15 S, with a resolution of 300. The buffer density, and the buffer viscosity were calculated by the program SEDNTERP (*43*), respectively. Figures of the sedimentation profile, c(s) distribution, and fitting results were generated using program GUSSI (*44*).

### Solid-state Multidimensional NMR

^13^C-^13^C DARR solid-state NMR experiments were performed by using ^13^C- and ^15^N-labelled αSyn fibrils prepared under the conditions of ionic strength and temperature indicated in the text. The ^13^C- and ^15^N-labelled αSyn was expressed in M9 minimal media containing ^13^C_6_-D-glucose and ^15^NH_4_Cl and purified as described for the unlabeled protein. The fibril samples were packed into 3.2 mm Varian-style solid-state NMR sample rotors (Phoenix NMR) using a home-made rotor packing tool (*45*) via ultra-centrifugation (92,000×g for two hours).

NMR spectra were acquired on a JEOL ECA-II 600 MHz NMR spectrometer equipped with a 3.2 mm Varian T3-HXY probe at a MAS rate of 12.5 kHz. A cooling gas was used to maintain the sample temperature to >20 ℃, estimated using the temperature-dependent longitudinal relaxation time of K^79^Br (*46*). The 90° pulse lengths of ^1^H and ^13^C were 3.2 μs. ^1^H to ^13^C cross-polarization (CP) was performed under a Hartman-Hahn matching condition of ∼50 kHz for ^1^H and ∼37.5 kHz for ^13^C (*47*), with a linear-ramp gradient applied to the ^13^C channel. ^13^C-^13^C mixing was achieved by 20 ms of DARR (*48*). During indirect and direct detection periods, ^1^H spins were decoupled by 70 kHz of SPINAL64 field (*49*). The acquisition time for both direct and indirect dimension was 10.24 ms with 10 μs of dwell time. The ^13^C chemical shift was referenced to sodium 2,2-dimethyl-2-silapentane-5-sulfonate (DSS) using the adamantane CH_2_ peak at 40.48 ppm (*50*).

All NMR data were processed with NMRPipe (*51*). The Lorentz-to-Gauss window function was applied to the data prior to the zero filling and Fourier transformation. The processed data were visualized by Python with the nmrglue (*46*) and Matplotlib (*52*) packages.

### Solution Multidimensional NMR

The uniformly ^15^N ([U-^15^N]) and ^13^C,^15^N ([U-^13^C,^15^N]) labeled αSyn was overexpressed in *E. coli* in M9 minimal medium containing ^15^NH_4_Cl and ^12^C-glucose or ^13^C-glucose as the solo nitrogen and carbon sources, respectively, and purified as described for the unlabeled protein. The NMR measurements were performed using an Avance III HD 800 spectrometer equipped with a TXI cryogenic probe and Avance III HD 600 spectrometer equipped with a QCI-P cryogenic probe (Bruker Biospin).

The following NMR experiments were performed at 37 °C for backbone signal assignment of αSyn; 3D HNCACB, HNCOCACB, HNCA, HNCOCA, HNCO, HNCACO, HNCANNH, and 2D ^1^H-^15^N HSQC. Additionally, chemical shift information for αSyn registered in the Biological Magnetic Resonance Data Bank was referenced (BMRB Entry 27074). The [U-^13^C,^15^N] labeled αSyn were dissolved in NMR buffer containing 50 mM TRIS-HCl buffer [pH 7.4], 150 mM NaCl, 2%(v/v) D_2_O. The concentrations of the αSyn were 100 µM.

^1^H-^15^N heterogenous single-quantum coherence (HSQC) experiments were performed using 100 µM [U-^15^N] labeled αSyn dissolved in fibrillation buffer prepared in H_2_O/D_2_O (98:2, v/v), under the ionic strength and temperature conditions specified in the text.

All NMR spectra were processed with Topspin (Bruker Biospin), NMRPipe (*51*) and POKY Suite (*53*).

### Circular dichroism (CD) spectroscopy

CD measurements were performed using a Jasco J820 spectrophotometer at 37 °C. Quartz cuvettes of 0.1 cm were utilized. The employed protein concentration was 0.25 mg mL^-1^ in fibrillation buffer. Spectra were recorded in 250-200 nm (far-UV CD) ranges and reported as the mean residue ellipticity ([θ]_MRW_, deg cm^2^ dmol^-1^) after subtracting the baseline.

### Ionic strength calculation

Ionic strength (IS) was calculated using the equation:

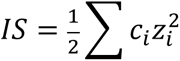

where *c*_i_ is the molar concentration of each individual ion, and *z*_i_ is the charge of each ion (*54*). In fibril formation and NMR experiments using Ca^2+^, where we fixed the ionic strength to 0.15 to mimic physiological environments, the ionic strength was set considering the necessary concentration of Ca^2+^ for each experiment and adjusted to 0.15 by adding the required amount of NaCl.

## Acknowledgments

We would like to thank Shizuka Sonoda and Miki Yoshida for protein expression and purification and the excellent technical support received. CA thanks JSPS (postdoctoral fellowship P16388) for the financial support.

## Funding

This work was supported by JSPS KAKENHI Grant JP24H00045, JSPS KAKENHI Grant JP22K07516 JSPS KAKENHI Grant JP22H02951 JSPS KAKENHI Grant JP22K15643 JSPS KAKENHI Grant JP23K24212 JSPS KAKENHI Grant JP23K18255

JSPS Core-to-Core Program A Advance Research Networks JPJSCCA20180007 JST FOREST Program JPMJFR231L

Japan Agency for Medical Research and Development (AMED) JP22gm1410014

Japan Agency for Medical Research and Development (AMED)JP24gm1910008

Japan Agency for Medical Research and Development (AMED)JP24ek0109771h

## Author contributions

Each author’s contribution(s) to the paper should be listed (we suggest following the CRediT model with each CRediT role given its own line. No punctuation in the initials.

Examples:

Conceptualization: CA, YM, MS, KK, YK, TO, KB, SN, YN, HO, SU, YM, KT, YG, KI and HM

Methodology: CA, YM, MS, HT, TM, JD, KY, KN, YY, HO, SU, YM, KT, YG, KI and HM

Investigation: CA, YM, MS, HT, TM, JD, NW, KY, KN, YY, HI, CJC, TA, YM, and KI

Supervision: YM, KK, YK, TO, KB, SN, YN, HO, SU, YM, KT, YG, KI and HM

Writing—original draft: CA and KI

Writing—review & editing: YM, KK, YK, TO, KB, SN, YN, HO, SU, YM, KT, YG, KI and HM

## Data and materials availability

All data is available in the main text or the supplementary materials.

