## Supplementary Materials for "Conformational switch in the alpha-synuclein C-terminus domain directs its fibril polymorphs"

### Affiliations

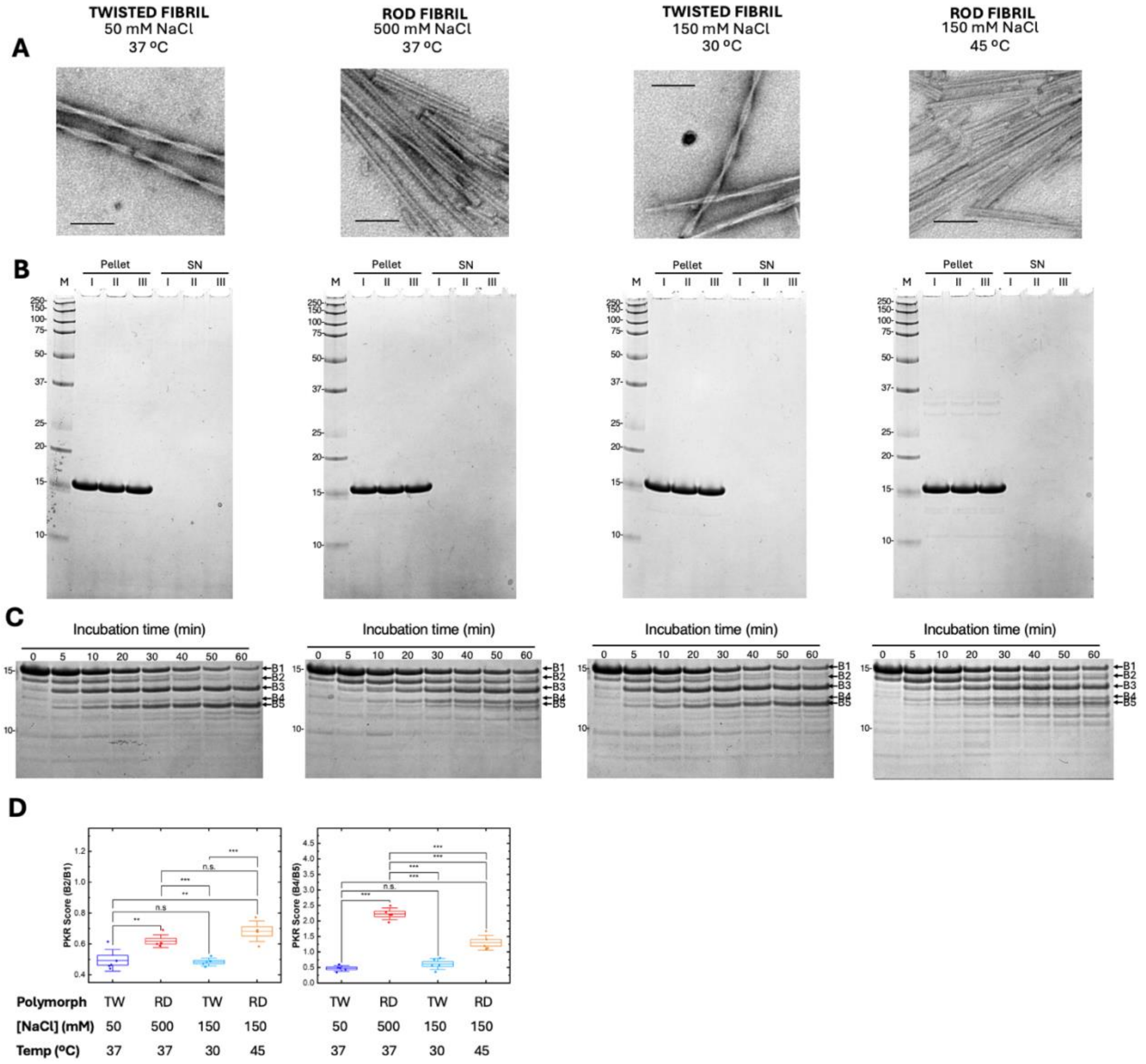


***Figure S1. Characterization of αSyn fibril morphologies. A****. Representative electron microscopy images of the analyzed fibrils (scale bar: 100 nm).* ***B.*** *Determination of the extent of the* ***A****. Representative TEM images of the analyzed fibrils (scale bar: 100 nm).* ***B.*** *Analysis of fibrillation completion under different conditions that generate twisted and rod-like polymorphs (conditions indicated in the upper legend). Fibril samples were ultracentrifuged, separated into the supernatants and the pellet fractions, and pellets were resuspended to the original volume of the indicated buffer before SDS-PAGE gel analysis. No detectable protein remained in the supernatant fractions. M: molecular weight markers; P: resuspended pellets; SN: supernatants.* ***C.*** *Time course of proteinase K digestion patterns for the twisted and rod-like fibrils shown in panel A. Analysis is focused on the five highest molecular weight bands (B1-B5). Molecular weight markers (kDa) are shown on the left of each panel.* ***D.*** *Densitometric analysis of 30-minutes digestion products showing intensity ratios B2/B1 and B4 /B5. TW: twisted fibril; RD: rod-like fibril. Boxes represent standard error of the mean values, while whiskers represent standard deviation. Statistical analyses were performed by Student's t-test (* $p < 0.05$; **$p < 0.01$; *** $p < 0.001$).*

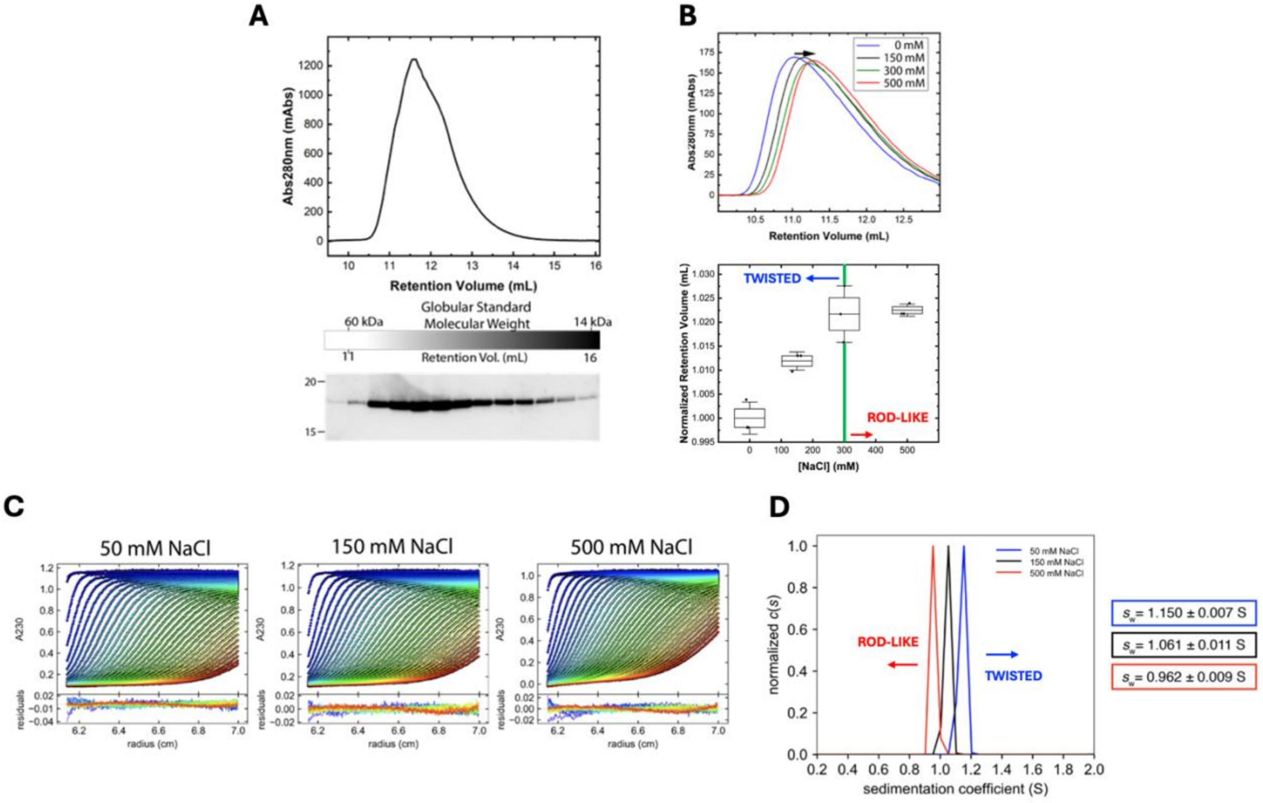


***Figure S2. Conformational diversity of αSyn monomer in solution. A.*** *A solution of αSyn in 50 mM TRIS buffer (pH 7.4) containing 300 mM NaCl was analyzed by SEC (upper) and subsequent Western Blotting of collected fractions using Anti-Syn211 antibody (lower).* ***B.*** *Upper: Representative SEC chromatograms of solutions of αSyn solution at different NaCl concentrations. Lower: Box-and-whisker plot of the normalized retention volumes. Boxes represent standard error of the mean values, whiskers represent standard deviation.* ***C.*** *Sedimentation Velocity Analytical Ultracentrifugation profiles of monomer αSyn in the presence of 50 mM (left), 150 mM (middle) and 500 mM NaCl (right). For clarity, data are shown at 18-minutes intervals.* ***D.*** *Apparent sedimentation coefficient values ($s_w$) of monomer αSyn were determined to be 1.150 ± 0.007 S, 1.061 ± 0.011 S and 0.962 ± 0.009 S at 50 mM, 150 mM and 500 mM NaCl, respectively. Values represent the average and standard deviation of three independent measurements (**Table S1**).*

***Table S1.*** *Apparent sedimentation coefficient values from the triplicated measurements, shown in Figure S2.D.*

| $s_w$ | 50 mM NaCl | 150 mM NaCl | 500 mM NaCl |
|---|---|---|---|
| Cell1 | 1.149 | 1.068 | 0.973 |
| Cell2 | 1.144 | 1.048 | 0.957 |
| Cell3 | 1.157 | 1.067 | 0.957 |
| Average | 1.150 | 1.061 | 0.962 |
| SD | 0.007 | 0.011 | 0.009 |

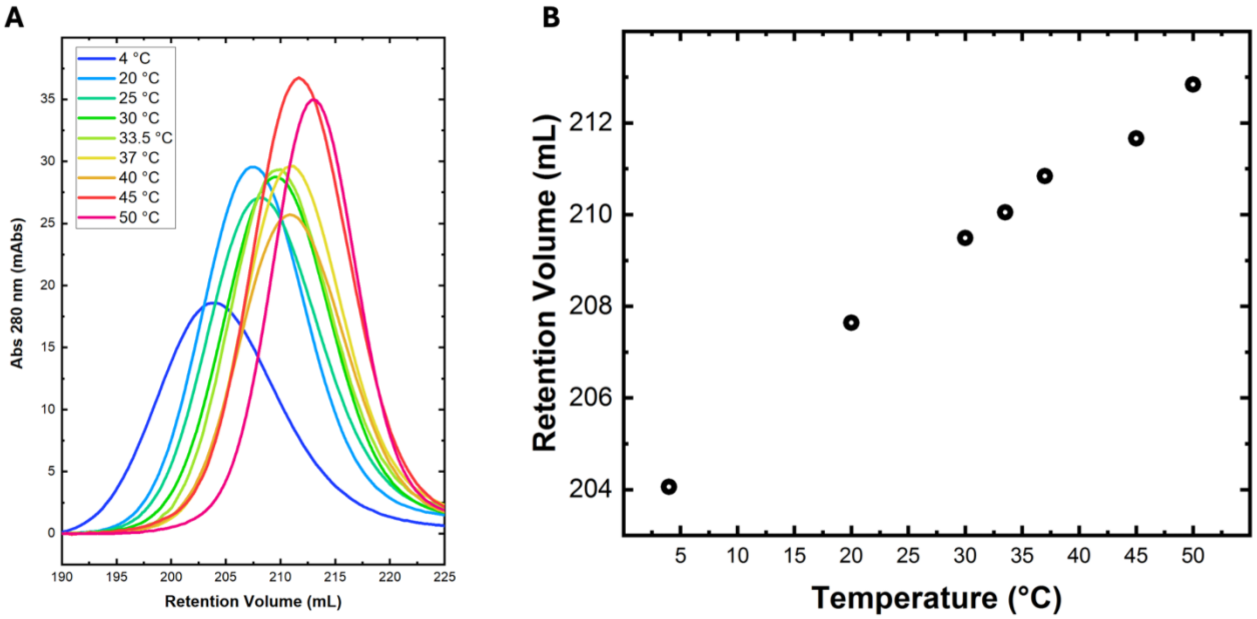


***Figure S3. SEC analysis of αSyn solutions at different temperatures. A.*** *SEC chromatograms of αSyn in 50 mM TRIS buffer (pH 7.4) containing 150 mM NaCl, measured at the indicated temperatures.* ***B.*** *Retention volume of monomeric αSyn as a function of temperature.*

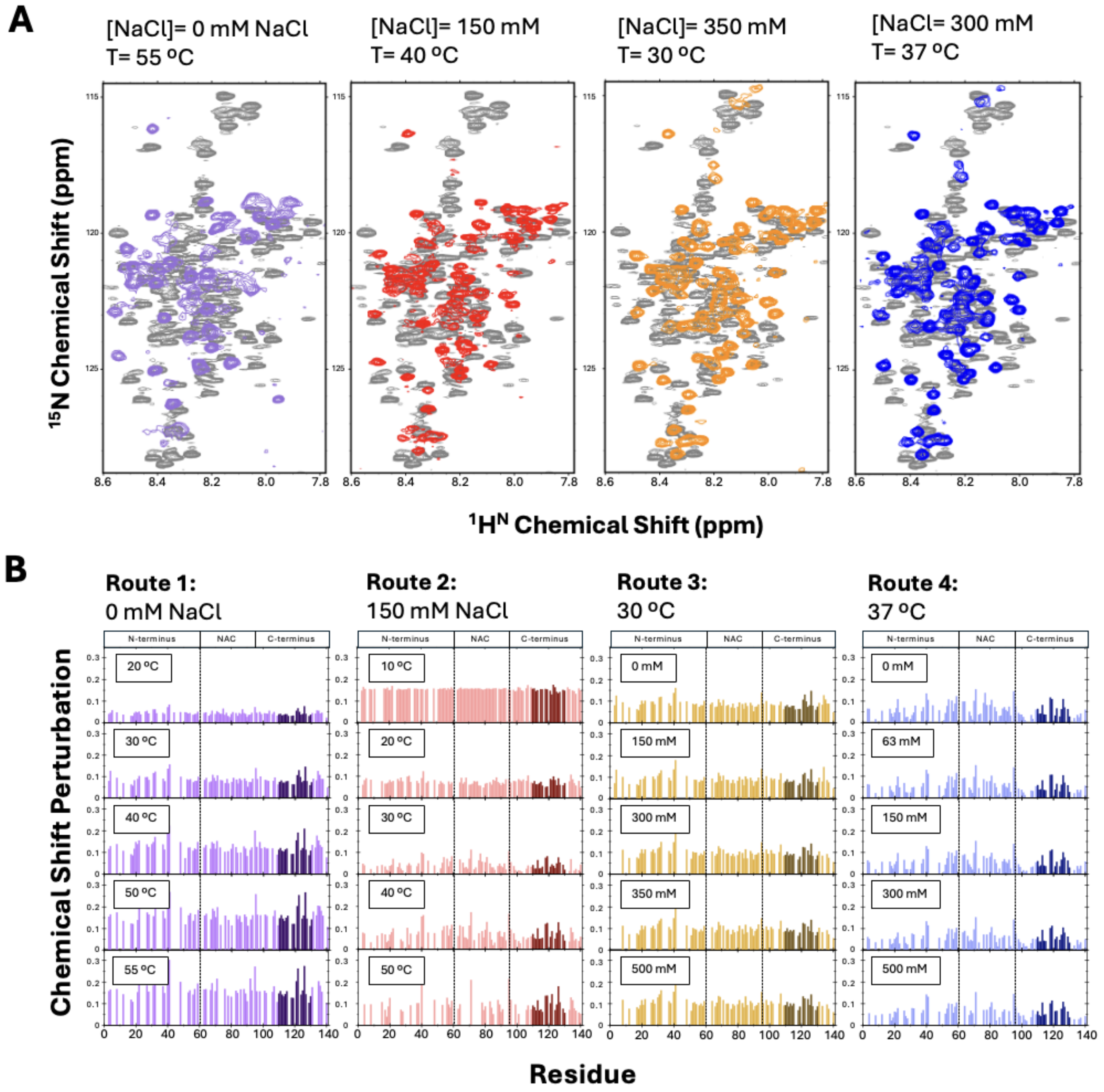


***Figure S4. Chemical shift perturbation across the routes presented in Figure 3.A. A.*** *Representative $^{1}$H-$^{15}$N HSQC spectra each route. The reference spectrum obtained at 10 ºC without NaCl is shown in gray.* ***B.*** *Chemical shift perturbation values for the $^{1}$H-$^{15}$N HSQC spectra of selected points along the routes R1 to R4, relative to the reference spectrum. The residues 110-130 are highlighted.*

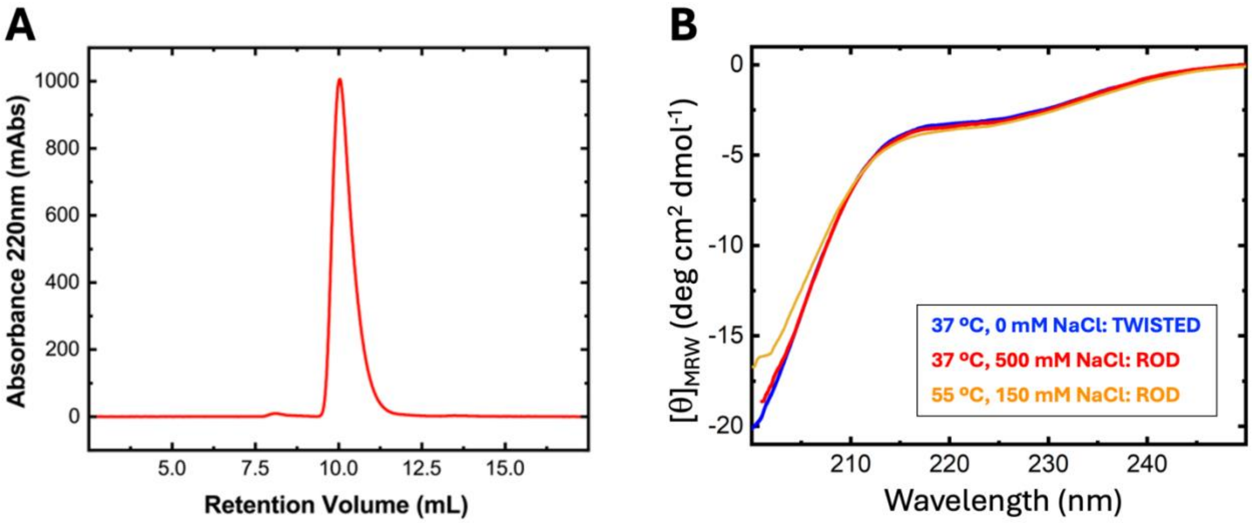


***Figure S5. $\alpha$Syn remains as a disordered monomer under the polymorph-inducing conditions. A.*** *SEC chromatograms of $\alpha$Syn in 50 mM TRIS buffer (pH 7.4) containing 500 mM NaCl after the $^{1}$H-$^{15}$N HSQC measurement at 37 ºC.* ***B.*** *Far-UV CD spectra of $\alpha$Syn solution under the polymorph-inducing conditions: i) 0 mM NaCl and 37 ºC for twisted fibrils (blue); ii) 500 mM NaCl and 37 ºC for rod fibrils (red); and 150 mM and 55 ºC for rod fibrils (yellow).*

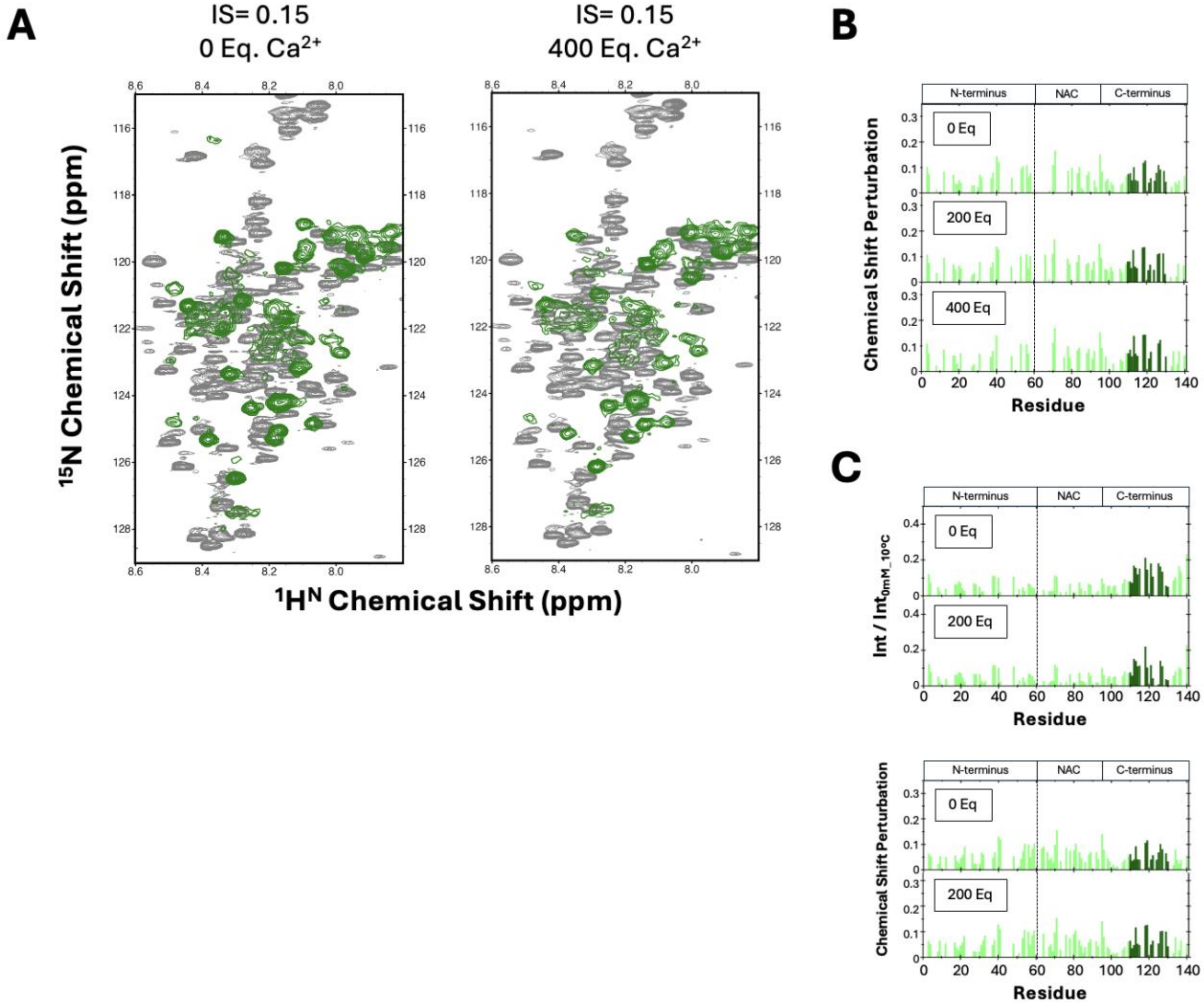


***Figure S6. NMR analysis on calcium-induced conformational change.*** *Overlay of $^{1}H$-$^{15}N$ HSQC spectra with and without 400 Eq. $Ca^{2+}$, referenced against the spectrum obtained at 10 ºC without NaCl (gray).* ***B.*** *Chemical shift perturbation at various $Ca^{2+}$ concentrations referenced to the spectrum obtained with no NaCl at 10 ºC. C-terminal residues (110-130) are highlighted.* ***C.*** *Upper: Peak intensity ratio from the $^{1}H$-$^{15}N$ HSQC spectra at different $Ca^{2+}$ concentrations (ionic strength of 0.2), normalized to the reference spectrum (no NaCl/$Ca^{2+}$ at 10 ºC). Lower: Chemical shift perturbation analysis at varying $Ca^{2+}$ concentrations referenced to the spectrum at 10 ºC without NaCl.*

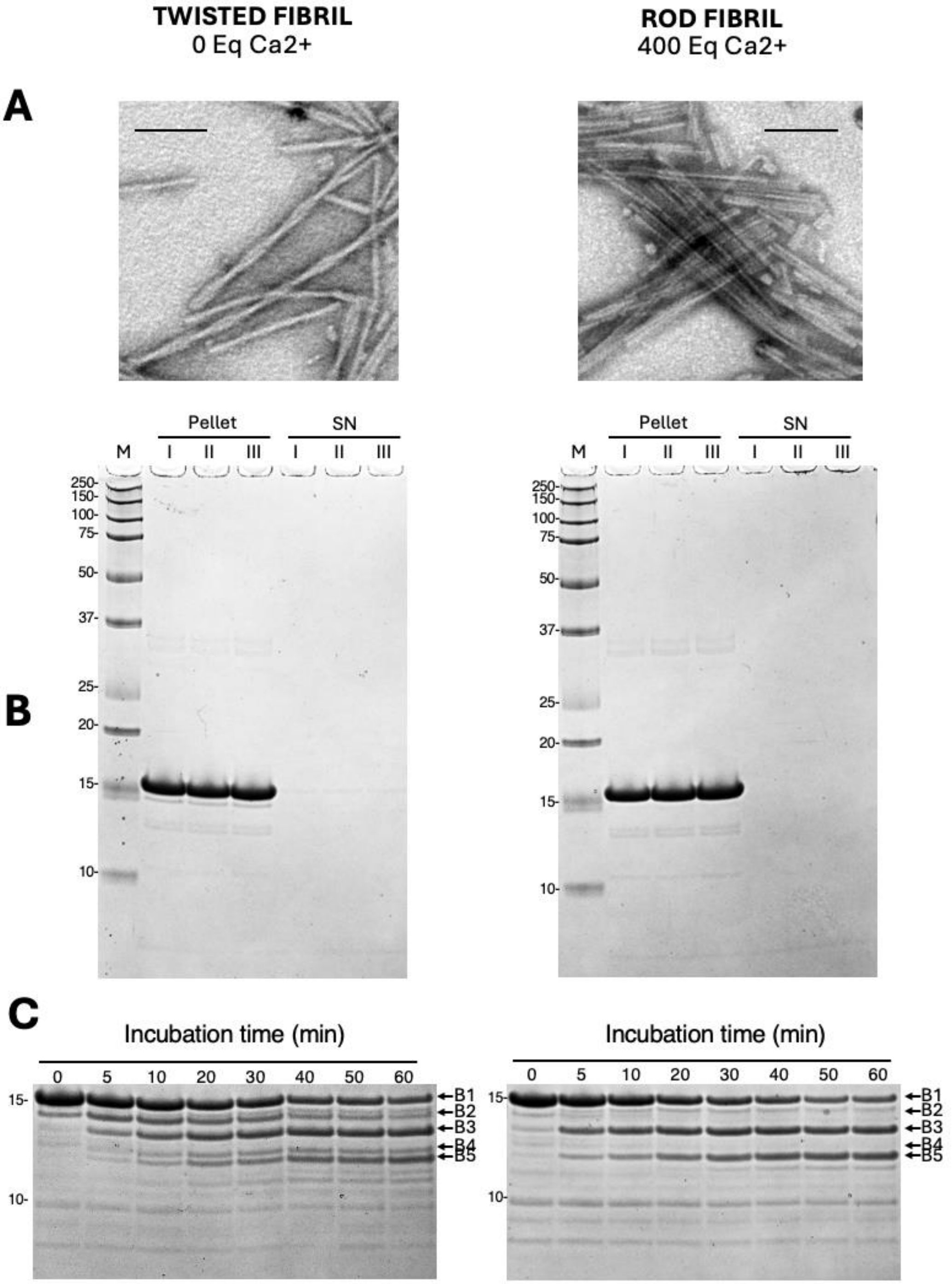


***Figure S7. Characterization of calcium-induced αSyn fibril polymorph.*** ***A***. *TEM images of fibrils produced in the presence and absence of calcium (scale bar: 100 nm).* ***B*** *Analysis of fibrillation completion in the presence or absence of calcium, at constant ionic strength (0.15). Fibril samples were treated as shown in Figure S1.A. In all cases, almost no detectable proteins were remained in the supernatant fractions. M: molecular weight markers; P: resuspended pellets; SN: supernatants*. ***C.*** *Time course of proteinase K digestion patterns for the fibrils in panel A. The first five highest molecular weight bands (B1-B5) were used to characterize the fragmentation patterns. Molecular weight markers (kDa) are shown on the left of each panel.*
